# Automated cell tracking using StarDist and TrackMate

**DOI:** 10.1101/2020.09.22.306233

**Authors:** Elnaz Fazeli, Nathan H. Roy, Gautier Follain, Romain F. Laine, Lucas von Chamier, Pekka E. Hänninen, John E. Eriksson, Jean-Yves Tinevez, Guillaume Jacquemet

**Affiliations:** Laboratory of Biophysics, Institute of Biomedicine, Faculty of Medicine, University of Turku, Turku, FI; Department of Pathology and Laboratory Medicine, Children’s Hospital of Philadelphia Research Institute, Philadelphia, PA 19104, USA; Turku Bioscience Centre, University of Turku and Åbo Akademi University, Turku, FI; Cell Biology, Faculty of Science and Engineering, Åbo Akademi University, Turku, Finland; MRC-Laboratory for Molecular Cell Biology, University College London, London, UK; The Francis Crick Institute, London, UK; Image Analysis Hub, C2RT, Institut Pasteur, Paris, FR

**Keywords:** Microscopy, Cell migration, Image analysis, StarDist, TrackMate, Deep-learning, Automated tracking

## Abstract

The ability of cells to migrate is a fundamental physiological process involved in embryonic development, tissue homeostasis, immune surveillance, and wound healing. Therefore, the mechanisms governing cellular locomotion have been under intense scrutiny over the last 50 years. One of the main tools of this scrutiny is live-cell quantitative imaging, where researchers image cells over time to study their migration and quantitatively analyze their dynamics by tracking them using the recorded images. Despite the availability of computational tools, manual tracking remains widely used among researchers due to the difficulty setting up robust automated cell tracking and large-scale analysis. Here we provide a detailed analysis pipeline illustrating how the deep learning network StarDist can be combined with the popular tracking software TrackMate to perform 2D automated cell tracking and provide fully quantitative readouts. Our proposed protocol is compatible with both fluorescent and widefield images. It only requires freely available and open-source software (ZeroCostDL4Mic and Fiji), and does not require any coding knowledge from the users, making it a versatile and powerful tool for the field. We demonstrate this pipeline’s usability by automatically tracking cancer cells and T cells using fluorescent and brightfield images. Importantly, we provide, as supplementary information, a detailed step-by-step protocol to allow researchers to implement it with their images.

---

The study of cell motility typically involves recording cell behavior, using live-cell imaging, and tracking their movement over time ^1,2^. To enable the analysis of such data, various software solutions have been developed ^3–9^. However, despite the availability of these computational tools, manual tracking remains widely used among researchers due to the difficulty in setting up fully automated cell tracking analysis pipelines. Automated tracking pipelines share a typical workflow that starts with a segmentation strategy that identifies the objects to track in each image. Tracking algorithms are then used to link these objects between frames. One challenging aspect of an automated tracking pipeline is often to achieve an accurate segmentation of the objects to track. One option to facilitate cell segmentation is to label their nuclei, using fluorescent dyes or protein markers. Nuclei can then be automatically segmented using intensity-based thresholding. However, this approach tends to become inaccurate when images are noisy or when the cells to track are very crowded ^10^. Deep-Learning approaches have demonstrated their robustness against these two issues ^11^. In this work, we present a new analysis workflow that builds upon a Deep-Learning segmentation tool and a cell tracking tool to achieve robust cell tracking in cell migration assays. We combine StarDist, a powerful deep learning-based segmentation tool, and Track-Mate, a user-friendly tracking tool, into a tracking pipeline that can be used without requiring expertise in or specialized hardware for computing (Figure 1) ^12–15^.

**Fig. 1.**
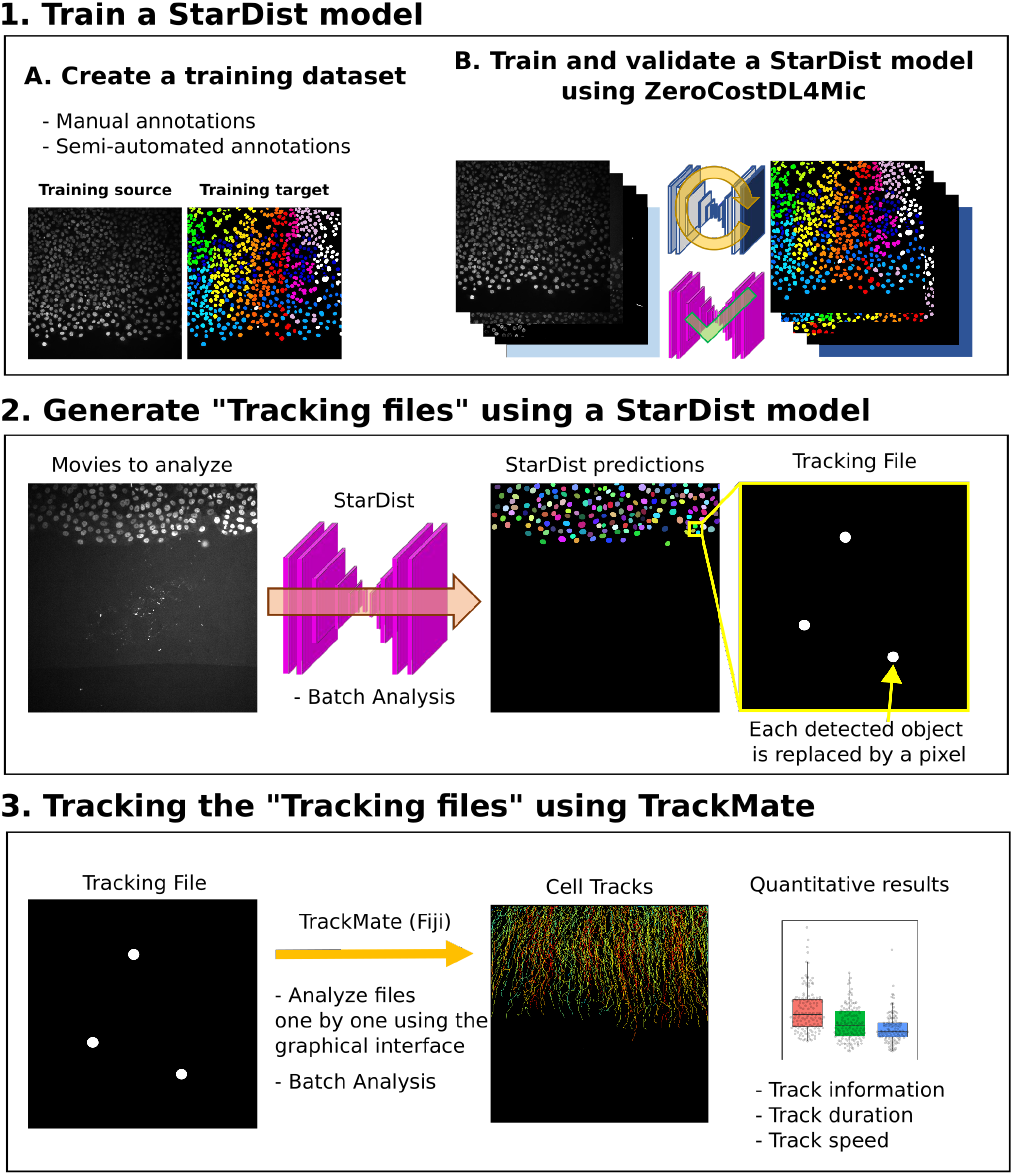
Workflow depicting how StarDist and TrackMate can be combined to track cells automatically.

The use of deep learning networks, such as StarDist, often requires the user to train or retrain a model using their images. While high-quality StarDist pre-trained models are readily available, they are likely to underperform when used on different data with, e.g., different staining, noise, and microscope type ^15^. To train StarDist models, we took advantage of the ZeroCostDL4Mic platform, allowing researchers to train (and retrain), validate, and use deep learning networks ^15^. Importantly, the ZeroCostDL4Mic StarDist 2D notebook can directly output a file containing all the nuclei’s geometric center coordinates (named tracking files), that can be used as input for TrackMate (Figure 1). Therefore, our proposed pipeline can be divided into three parts (Figure 1; Supplementary protocol).

- First, a StarDist model is trained using the Zero-CostDL4Mic platform. This part needs to be performed only once for each type of data.
- Second, the trained StarDist model is used to segment the object to track and generate Tracking files.
- Finally, the tracking files can be used in TrackMate to track the identified objects.

Training a StarDist model requires a set of images and their corresponding masks (Figure 1 and Figure 2). Generating a training dataset is by far the most time-consuming part of the analysis pipeline presented here as it requires the manual annotations of the images to analyze (Supplementary protocol). For instance, to generate the training datasets presented in Figure 2, each cell/nuclei contour was drawn manually using the freehands selection tool in Fiji. The creation of a high-quality training dataset is a critical part of the process as it will impact the specificity and performance of the StarDist model. However, the generation of a training dataset is only required once per dataset type. If a StarDist model already exists for similar images it can be used to significantly accelerate the creation of the training dataset via semi-automated annotation (see Supplementary protocol).

**Fig. 2.**
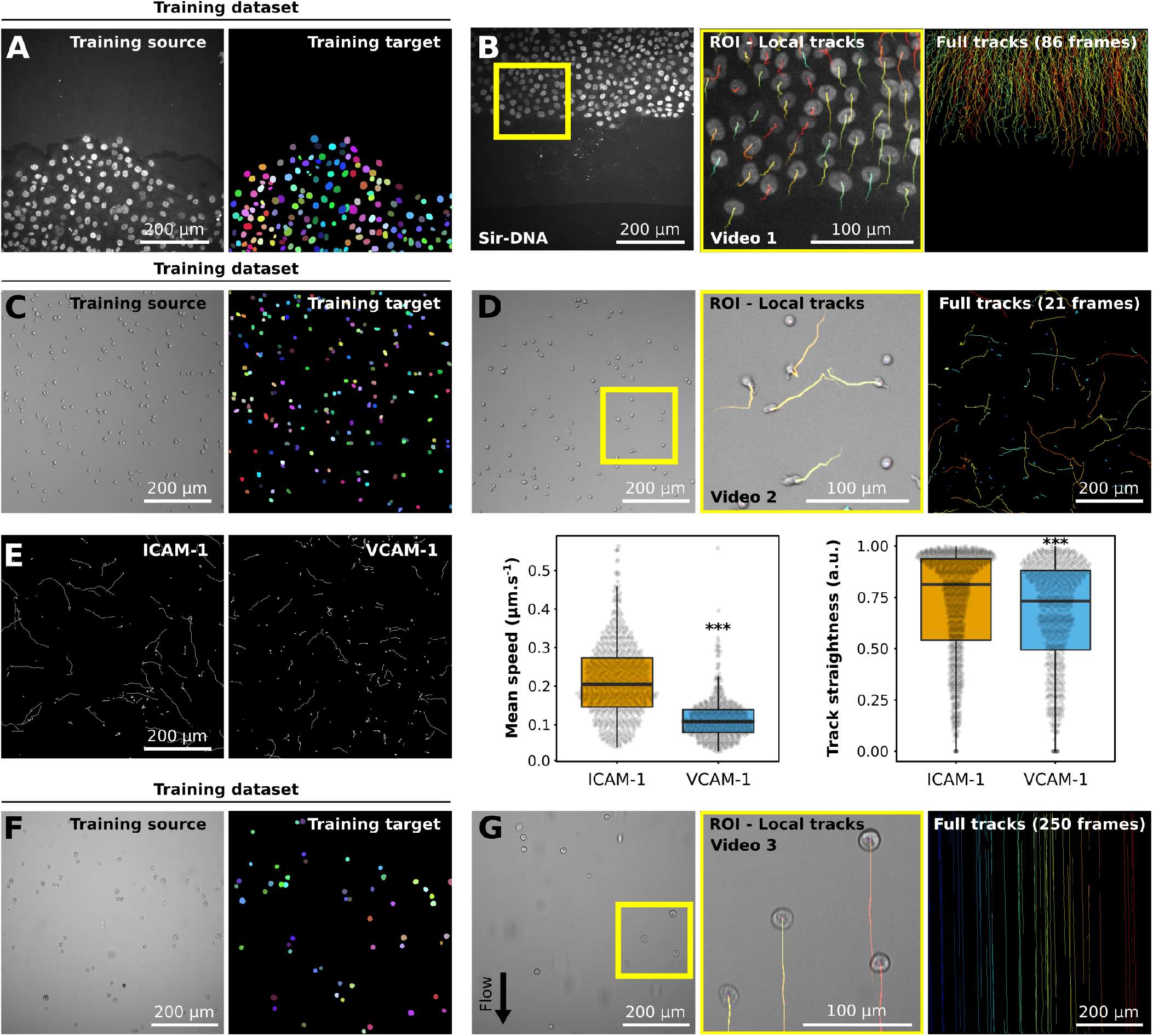
Example of datasets analyzed using StarDist and TrackMate. **A-B**: Migration of MCF10DCIS.com, labeled with Sir-DNA, recorded using a spinning disk confocal microscope and automatically tracked. Examples of images used to train StarDist (**A**), and an example of results obtained using automated tracking are displayed (**B**, Video 1). The yellow square indicates a magnified ROI, where the local track of each nucleus is displayed. The full cell tracks are displayed on the left. Tracks are color-coded as a function of their maximum instantaneous velocity (blue slow, red fast tracks). **C-E**: Migration of activated T cell plated on VCAM-1 or ICAM-1, recorded using a brightfield microscope and automatically tracked. Examples of images used to train StarDist (**C**) and an example of results obtained using automated tracking are displayed (**D**, Video 2). (**E**) Comparison of the migration of activated T cells on VCAM-1 or ICAM-1. Track mean speed and track straightness were quantified. Data are displayed as boxplots17. *** p-value = <0.001, p-values were determined using a randomization test. **F-G**: Cancer cells flowing in a microfluidic chamber, recorded live using a brightfield microscope and automatically tracked (Video 3). Examples of images used to train StarDist (**F**), and an example of results obtained using automated tracking are displayed (**G**). The full tracks shown here were color-coded as a function of their x coordinate.

One of our analysis pipeline’s key features is that, once a StarDist model has been satisfactorily trained, movies of migrating cells can efficiently be processed in batch. Indeed, while individual tracking files can be analyzed one by one using the TrackMate graphical interface, we also provide a Fiji macro to analyze a folder containing multiple tracking files. Our batch processing macro will provide basic quantitative information for each track, including median and maximal speeds. If more information is needed, the tracking results generated by our script are directly compatible with the Motility lab website, where they can be further processed ^16^.

To illustrate our analysis pipeline’s functionality and flexibility, we first trained a StarDist model to analyze the behavior of breast cancer cells migrating collectively (Figure 2A, Video 1). The cancer cell’s nuclei were fluorescently labeled, and the cells imaged using fluorescence-based microscopy. The creation of the training dataset used in this example was greatly facilitated by the availability of a StarDist model, released by the StarDist creators, capable of segmenting fluorescent nuclei. In this case, the StarDist Fiji plugin was used to segment the location of nuclei in the training images, and all miss-annotations were manually corrected (Supplementary protocol). To highlight that our pipeline can also be used to analyze brightfield images, we generated a StarDist model to track T cells migrating on ICAM-1 or VCAM-1 (Figure 2C-E, Video 2). Importantly, automated analysis of these data could reproduce the results obtained via manual tracking ^17^. Finally, we used our pipeline to automatically track non-adherent cancer cells flowing in a microfluidic chamber (Figure 2F and 2G, Video 3). In this case, automated tracking is especially useful due to the very high number of frames to analyze. For the last two examples, no suitable pretrained StarDist models were available. Therefore, to generate the training datasets, we manually annotated 20 images and trained a first StarDist model. This model was then used to accelerate the annotation of the rest of the training images.

Here we show that StarDist and TrackMate can be integrated seamlessly and robustly to automate cell tracking in fluorescence and brightfield images. We envision that this pipeline can also be applied to any circular or oval-shaped objects. However, we acknowledge that using brightfield images may not always work directly with our pipeline, especially if cells display complex and interchanging shapes since StarDist is mostly designed to detect round or compact shapes. In this case, other tools, such as Usiigaci, could also be considered ^8^. Still, brightfield images could also be artificially labeled using deep learning, transforming the brightfield dataset into a pseudo-fluorescence one, as can be done with ZeroCostDL4Mic already ^15^. The pipeline described here is currently limited to the tracking of objects in 2D. However, a similar workflow can be applied to 3D datasets as both StarDist and TrackMate can accommodate 3D images ^12,13,18^.

## Material and methods

### Breast cancer cell dataset ^15^

MCF10DCIS.com cells were described previously^19^. DCIS.COM lifeact-RFP cells were incubated for 2h with 0.5 µM SiR-DNA (SiR-Hoechst, Tetubio, Cat Number: SC007) before being imaged live for 14h using a spinning-disk confocal microscope (1 picture every 10 min). The spinning-disk confocal microscope used was a Marianas spinning disk imaging system with a Yokogawa CSU-W1 scanning unit on an inverted Zeiss Axio Observer Z1 microscope (Intelligent Imaging Innovations, Inc.) equipped with a 20x (NA 0.8) air, Plan Apochromat objective (Zeiss).

### T cell dataset ^17^

Lab-Tek 8 chamber slides (ThermoFisher) were coated with 2 µg/mL ICAM-1 or VCAM-1 overnight at 4°C. Activated primary mouse CD4+ T cells were washed and resuspended in L-15 media containing 2 mg/mL D-glucose. T cells were then added to the chambers, incubated 20 min, gently washed to remove all unbound cells, and imaged. Imaging was done using a 10x phase contrast objective at 37°C on a Zeiss Axiovert 200M microscope equipped with an automated X-Y stage and a Roper EMCCD camera. Time-lapse images were collected every 30 sec for 10 min using SlideBook 6 software (Intelligent Imaging Innovations).

### Flow chamber dataset

AsPC1 pancreatic cancer cells (500 000 cells/ml in PBS) were perfused at a speed of 300 µm/sec using a peristaltic pump (ISMATEC MS12/4 analogic) and a homemade tubing system (Ismatek 3-Stop tubes and Ibidi® tubings and connectors) in a microchannel (Ibidi® µ-slides400 LUER). Images were acquired with a brightfield microscope (Zeiss Laser-TIRF 3 Imaging System, Carl Zeiss) and a 10X objective.

### Data display and statistical analyses

Box plots were generated using PlotsOfData ^20^. Randomization tests were performed using the online tool PlotsOfDifferences ^21^.

### Availability

All required software is fully open-source and can be accessed on GitHub. The training datasets used to train StarDist, example images and the trained StarDist models are available for download in Zenodo (Breast cancer cell dataset, T cell dataset, Flow chamber dataset).

## Supplementary video

**Video 1: Automated tracking of breast cancer cell migrating collectively**. MCF10DCIS.com cells, labeled with Sir-DNA, were recorded using a spinning disk confocal microscope and automatically tracked using StarDist and Track-Mate. Local tracks are displayed.

**Video 2: Automated tracking of T cell migrating on ICAM-1**. Activated T cell plated ICAM-1 were recorded using a brightfield microscope and automatically tracked using StarDist and TrackMate. Local tracks are displayed.

**Video 3: Automated tracking of cancer cells flowing in a microfluidic chamber**. AsPC1 pancreatic cancer cells flowing in a microfluidic chamber were recorded live using a brightfield microscope and automatically tracked using StarDist and TrackMate. Local tracks are displayed.

## ACKNOWLEDGEMENTS

This work was supported by grants awarded by the Academy of Finland (G.J.), the Sigrid Juselius Foundation (G.J.), Åbo Akademi University Research Foundation (CoE CellMech; G.J.) and by Drug Discovery and Diagnostics strategic funding to Åbo Akademi University (G.J.). G.F. was supported by the National Cancer Center Finland (FICAN). NHR acknowledges support from the Cancer Research Institute and NIH (T32 AR007442). R.F.L. acknowledges the support of the MRC Skills development fellowship (MR/T027924/1).

The Cell Imaging and Cytometry Core facility (Turku Bioscience, University of Turku, Åbo Akademi University, and Biocenter Finland), the Finnish Euro-Bioimaging Node, and Turku Bioimaging are acknowledged for services, instrumentation and expertise.

## AUTHOR CONTRIBUTIONS

G.J. conceived the project; E.F., L.v.C., R.F.L., J.Y.T., and G.J. wrote the code; N.H.R, G.F., and G.J. performed the image acquisition of the example data; E.F. and G.J. wrote the manuscript with input from all co-authors.

## DETAILED PROTOCOL

### EQUIPMENT/ EQUIPMENT SETUP

- A computer with access to the internet.
- A Google account. You can create a free account here.
- The ZeroCostDL4mic StarDist 2D notebook. The latest version of this notebook is available here.
- A Fiji script to batch analyze movies. The latest version of this script is available here.
- The latest version of Fiji. TrackMate and its dependencies are already installed in Fiji. You can download Fiji here.
- The Laboratory for Optical and Computational Instrumentation (LOCI) Fiji plugin
- To install the LOCI plugin, In Fiji, click on [Help > Update]. Then click on Manage update sites. Find “LOCI” from the list and tick the box. Close the Manage update sites window. In the Fiji Updater window, click on Apply changes. Restart Fiji.
- The StarDist Fiji plugin. To install the StarDist plugin, In Fiji, click on [Help > Update]. Then click on Manage update sites. Find “StarDist” from the list and tick the box. Close the Manage update sites window. In the Fiji Updater window, click on Apply changes. Restart Fiji.
- Images to analyze.

## PROCEDURE

Before starting your analysis, we recommend that you familiarise yourself with the StarDist, TrackMate and ZeroCostDL4Mic paper.

### Part 1: Train a StarDist network

Part 1 is done only once per dataset type (combination of microscopy type, magnification etc.). Once a suitable StarDist model is available, only Parts 2 and 3 need repeating.

#### 1.1: Generate a training dataset for StarDist

To train a StarDist model, you need matching images of nuclei/or cells and their corresponding masks. These images need to have the same name and to be saved in two different folders. The mask images can be generated in Fiji, either manually or semi-automatically. The semi-automatic approach can be used if an existing StarDist model performs reasonably well on your images. Alternatively, you can manually annotate 20 or so images, train your StarDist model, and then use the semi-automated annotation method to increase your training dataset size rapidly.

### Alternative A: Manual annotations using Fiji and the LOCI plugin

- Open one image in Fiji.
- Open the ROI manager [Analyze-> Tools -> ROI Manager].
- Using the freehand selection tool, draw outlines around each nucleus manually.
- Add each outline to the ROI manager (key shortcut “t”).
- Once all the nuclei outlines are stored in the ROI manager, use the LOCI plugin to create an ROI map [Plugins-> LOCI-> ROI map]. This is your mask image.
- Rename the mask image so that its name matches the corresponding nuclei image and save it inside another folder.

### Alternative B: Semi-automated annotations using the StarDist and LOCI Fiji plugins

- Open one image in Fiji.
- Open the StarDist Fiji plugin [Plugins-> StarDist> StarDist 2D].
- Choose the Model you want to use. If you use a model, you trained yourself (ie using ZeroCostDL4Mic), choose “Model from File,” and indicate the model path.
- Select “ROI Manager” as an Output Type.
- Press OK.
- From the ROI manager, check that each ROI corresponds to a nuclei. Delete each misidentification and manually add all the missing nuclei.
- Once all the nuclei outlines are stored in the ROI manager, use the LOCI plugin to create an ROI map (Plugins-> LOCI-> ROI map). This is your mask image.
- Rename the mask image so that its name matches the corresponding nuclei image and save it inside another folder.

#### 1.2: Train StarDist using the StarDist 2D ZeroCostDL4Mic notebook

In this section, you will find some information on how to use the ZeroCostDL4Mic StarDist 2D notebook. More detailed information on using ZeroCostDL4Mic notebooks can be found in the ZeroCostDL4Mic wiki, manual or explanatory videos. The output of this step is a trained StarDist model that can then be used on new data.

##### 1.2.1 Prepare your files for ZeroCostDL4Mic

When using deep learning networks, it is critical to be able to assess their performance. This is typically done by comparing the prediction generated by the model to ground truth images. This essential quality control (QC) step can be done directly in the ZeroCostDL4Mic notebook. Therefore, we recommend that you split the training dataset you generated into “training” and “QC” dataset. Your QC dataset can be composed of 2-4 images.

- Split your annotated images into “training dataset” and “QC dataset.”
- Upload your images to google drive.
- Open the StarDist (2D) notebook from the following link by clicking the “Open in Colab” icon.
- Create a copy of the notebook in your GDrive. [File> Save a Copy in Drive].

##### 1.2.2 Initialise the ZeroCostDL4Mic notebook

In google Colab, to run a cell, drag the mouse cursor to the brackets [] and press the play button. After executing the code, the play icon is replaced with a number. Run each cell one after another. In the StarDist (2D) notebook, sections 1 and 2 allow you to initialize the notebook while sections 3 and 4 will enable you to train your StarDist model. Section 5 will allow you to assess the quality of your model, while section 6 will allow you to make predictions and generate tracking files.

- Mount your GDrive
- Install StarDist and the required dependencies

##### 1.2.3 Start Training

- Set the main parameters required to train StarDist (section 3.1 of the notebook).
- Locate your training dataset. The “training source” is the path to the folder containing your raw images, while the “training target” is the folder’s path containing your masks. There are three icons on the left side of the Google Colab page. Click on the folder icon to navigate your google drive and reach the folders of interest, right-click on the folder and copy path. Paste the path to each section to fill the parameters.
- Choose a name for your model and where to save it.
- Choose the number of EPOCH to use to train StarDist. We recommend that you train new models for at least 100-200 EPOCH.
- Run the cell.
- Optional - Use data augmentation to increase the size of your training dataset artificially (Section 3.2). Use this section only If your training dataset is small.
- Optional - Use Transfer Learning. If a pre-existing StarDist model has been trained on data that are somewhat similar to yours and performs relatively well on your data, consider using it as a starting point for training. This will accelerate the training process and can improve overall performance.
- Run section 4.1 to prepare your images for training
- Run section 4.2 to Start training the network. This step can take several hours to complete. At the end of the training, the model will be automatically saved in GDrive. In addition, the model will be automatically exported so it can be used to make predictions in the StarDist Fiji plugin.
- Download the trained StarDist model so it can be used in the StarDist Fiji plugin. Automated tracking is, however not supported via the Fiji StarDist plugin.

#### 1.3: Evaluate a StarDist model

Section 5 of the ZeroCostDL4Mic notebook allows you to assess the quality and performance of your trained model.

- Select the model to assess If you are assessing a model trained in a previous session, remember to initialize the Zero-CostDL4Mic notebook by running sections 1 and 2.
- Inspect the training curves (more information in the notebook) Compare the predictions made by your model to ground truth segmentations (more information in the notebook).

### Part 2: Generate Tracking files using a StarDist model

Section 6 of the StarDist ZeroCostDL4Mic notebook allows you to make predictions on your data and to generate Tracking files. If you are using a model that was trained in a previous session, remember to initialize the notebook by running sections 1 and 2.

- Input the location of your “Data_folder.”
- Choose the location of your “Results_folder.”
- Choose “Stacks” as “Data_type”
- Choose the type of generated outputs. In this workflow, Tracking_files are needed.
- Run the cell
- Once the predictions are completed, download the generated tracking files and other StarDist results.

### Part 3: Automated tracking using TrackMate

### Alternative A: Analyze your tracking files one by one using the TrackMate graphical interface

- Run Fiji
- Open a tracking file.
- Open TrackMate [Plugins > Tracking > TrackMate].
- If Fiji interprets the third dimension as z and not T, a new window will pop up to swap these two parameters. In that case, click “Yes.”
- Once Trackmate is open, click Next.
- Choose the Downsample LoG detector.
- Use the following parameters: Estimated blob diameter = 1, Threshold = 1 and downsampling factor = 1. Use the preview to ensure that the detection is correct.
- After the detection step, click on “Next” twice.
- Choose HyperStack Displayer and click on “Next” twice.
- Choose the LAP tracker.
- Choose an appropriate frame linking distance.
- If your cells are expected to divide, enable “Track segment splitting”; otherwise, disable it. Disable “Track segment merging.”
- After displaying the tracks and checking that the set distance parameters worked, save the tracking file by clicking on the Save button from the lower left part of the TrackMate window.
- To display statistical information on the cell tracks (such as track duration or track speed), click on analysis. Results are now ready to be further analyzed.
- To generate an image of the tracks, choose the appropriate parameters for Track display mode and the Set color by options, and then click Next. At the following step, click Next. Then click Execute.

### Alternative B: Batch process your tracking files

We recommend that you perform the analysis with the graphical interface once before performing a batch analysis. This will help you identify the best parameters.

- Download the required Fiji macro here.
- Open the macro in Fiji and press Run.
- Fill in the parameters based on your experimental parameters. Pixel and time calibration
- Choose an appropriate frame linking distance and enable/disable “Track segment splitting” and “Track segment merging.”
- Select the input directory containing the tracking files to analyze.
- Select where to save the results generated by TrackMate.
- This macro will generate two CSV files for each tracking file. One will contain some necessary information about each track, including their maximal and mean speed. The other file will contain the coordinate of each identified cell center and the track they belong to. This file can be further analyzed using the Motility lab website Motility lab website.

